# Direct visualization of how Actin Depolymerizing Factor’s filament severing and depolymerization synergizes with Capping Protein's "monomer funneling" to promote rapid polarized growth of actin filaments

**DOI:** 10.1101/114199

**Authors:** Shashank Shekhar, Marie-France Carlier

## Abstract

A living cell’s ability to assemble actin filaments in intracellular motile processes is directly dependent on the availability of polymerizable actin monomers which feed polarized filament growth. Continued generation of the monomer pool by filament disassembly is therefore crucial. Disassemblers like ADF/cofilin and filament cappers like Capping Protein (CP) are essential agonists of motility, but the exact molecular mechanisms by which they accelerate actin polymerization at the leading edge and filament turnover has been debated for over two decades. While filament fragmentation by ADF/cofilin has long been demonstrated by TIRF, filament depolymerization was only inferred from bulk solution assays. Using microfluidics-assisted TIRF microscopy, we provide the first direct visual evidence of ADF's simultaneous severing and rapid depolymerization of individual filaments. We have also built a conceptually novel assay to directly visualize ADF’s effect on a filament population. We demonstrate that ADF’s enhanced pointed-end depolymerization leads to an increase in polymerizable actin monomers co-existing with filaments, thus promoting faster barbed-end growth. We further reveal how ADF-enhanced filament depolymerization synergizes with CP’s long-predicted “monomer funneling” and leads to skyrocketing of filament growth rates, close to estimated rates in the lamellipodia. The “Funneling model” hypothesized, on thermodynamic grounds, that at high enough extent of capping, the few noncapped filaments transiently grow much faster, an effect proposed to be very important for motility. We provide the first direct microscopic evidence of monomer funneling by CP at the scale of individual filaments. We believe that these results enlighten our understanding of the turnover of cellular actin networks.

## Introduction

Force is produced against membranes by filament barbed-end assembly, which requires maintenance of a high concentration of polymerizable actin monomers[1, 2]. The monomer pool feeds the growing barbed ends, and is maintained at a steady state level by pointed-end filament depolymerization. In this treadmilling concept, the average rate of barbed-end growth over the filament population balances the rate of pointed-end depolymerization. Since pure actin filaments treadmill very slowly [3] compared to the rapid apparent treadmilling rates observed in cellular actin arrays (e.g. in extending lamellipodia) cellular upregulation of the intrinsically slow treadmilling is required [4].

ADF/cofilin and Capping Protein are known to play a key role in motility *in vivo* [5-9] as well as in reconstituted *in vitro* assays[10-12]. However, the molecular mechanisms underlying how ADF enhances motility (in absence of Capping protein) is still unclear [13-16]. Majority of studies on single filaments thus far have found ADF/cofilin to only fragment actin filaments [17-20]. However, severing (in absence of Cappers) will not promote an increase in the steady-state monomer concentration. Each of the new pointed ends created by severing will still depolymerize at their inherent slow rate of depolymerization (0.2 su/s), thus not enhancing the monomer pool and thereby maintaining identical slow treadmilling of single filaments [1]. One study however has proposed that ADF-mediated enhancement of filament depolymerization, measured via bulk solution light scattering experiments, was responsible for ADF’s effect on motility[21–23]. This proposal has been a debated issue since ADF’s enhanced depolymerization of actin filaments has so far never been microscopically visualized. Disassembly by ADF has been attributed to shortening by fragmentation [20]. In the present study, we provide the first visual evidence of ADF’s filament depolymerization in addition to well-characterized severing activities.

Capping proteins has also been shown to enhance motility in a concentration dependent fashion [10, 24–26]. When a large number of filaments are exposed to capping protein, the fraction of filaments getting capped will be dependent on CP concentration (fraction capped, 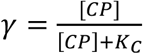, where Kc =0.1nM, is the equilibrium dissociation constant for CP binding to barbed ends). When an increasing fraction of filaments get capped, monomers released from all pointed ends feed the growth of only a few barbed ends that are not capped. Since these few noncapped barbed ends are fed from depolymerization of 100% of pointed ends, they individually grow much faster than if all barbed ends were free to grow (a “funneling effect”, Figure 4b) [27]. Another study has contested this model, and instead proposed that Capping Protein enhances nucleation [16].

So far, the effects of ADF’s enhanced depolymerization of filaments and CP’s funneling effect have only been interpreted from bulk solution experiments. Nevertheless, ADF in association with its effector Aip1 is known to cause enhanced single filament disassembly [28, 29]. Here we first use microfluidics-assisted single filament imaging [30, 31] to reveal both severing and rapid depolymerization of actin filaments by ADF alone. This assay is then adapted to monitor the effects of ADF and Capping Protein (CP) (alone and together) on the concentration of ATP-monomers co-existing with filaments at steady state, thus providing direct visual evidence of funneling effect. Filaments anchored at one of their ends can be exposed to a flow of known biochemical composition. Flow doesn’t alter kinetics of protein binding[32]. The key advantage is that filaments do not need to be tethered to the surface via biotin/myosin linkers, which can interfere with the actin twist and affect filament interactions with side-binding proteins like ADF. Results presented here provide the first single-filament demonstration of how ADF and CP work synergistically to enhance treadmilling at the level of individual filaments.

## Results and Discussion

### ADF increases the rate of depolymerization at pointed and barbed ends

We first studied the effect of ADF on filament pointed ends. Filaments initiated from surface-anchored gelsolin-actin complexes were allowed to elongate at pointed ends. Filaments were then aged for 15 minutes to convert to ADP-F-actin. These filaments were then exposed to a flow of polymerization buffer (F-buffer: 5 mM Tris-Cl-pH 7.8, 2 mM ATP, 1 mM MgCl2, 0.2 mM EGTA, 50 mM KCl, 10 mM DTT, and 1 mM DABCO) containing increasing amounts of ADF (Figure 1a). Filaments exposed to F-buffer alone expectedly did not appreciably fragment within the observation period of 10 min and their pointed ends depolymerized extremely slowly (0.17±0.04 subunits/s, S.D; n=50 filaments, 3 independent experiments), in agreement with earlier single filament studies (0.16 su/s) [33]. These values are lower than the ones derived from bulk solution measurements [3], possibly due to the slower dissociation of photo-induced F-actin dimers [30]. Upon exposure to ADF, we observed both severing of filaments and faster pointed-end depolymerization between consecutive severing events (Figure 1b). The frequency of severing increased with ADF concentration, as seen previously [18, 34]. Figure 1e shows that 50% of filaments (mean length = 10μm) have severed once at t_1/2_ = 40s in the presence of 5 μM ADF, consistent with the scheme F -> 2F and a first order fragmentation rate constant of ln(2)/t_1/2_ = 0.017 s^-1^ for a 10 μm long filament. The rate of pointed-end depolymerization also increased with ADF concentration in a saturating fashion up to a maximal rate of 3.72±0.35 su/s (n=50 filaments, 3 independent experiments) at 5 μM ADF (Figure 1c). The 22-fold increase in depolymerization rate at pointed ends is similar to the 25fold increase observed in bulk solution measurements [21].

**Figure 1:**
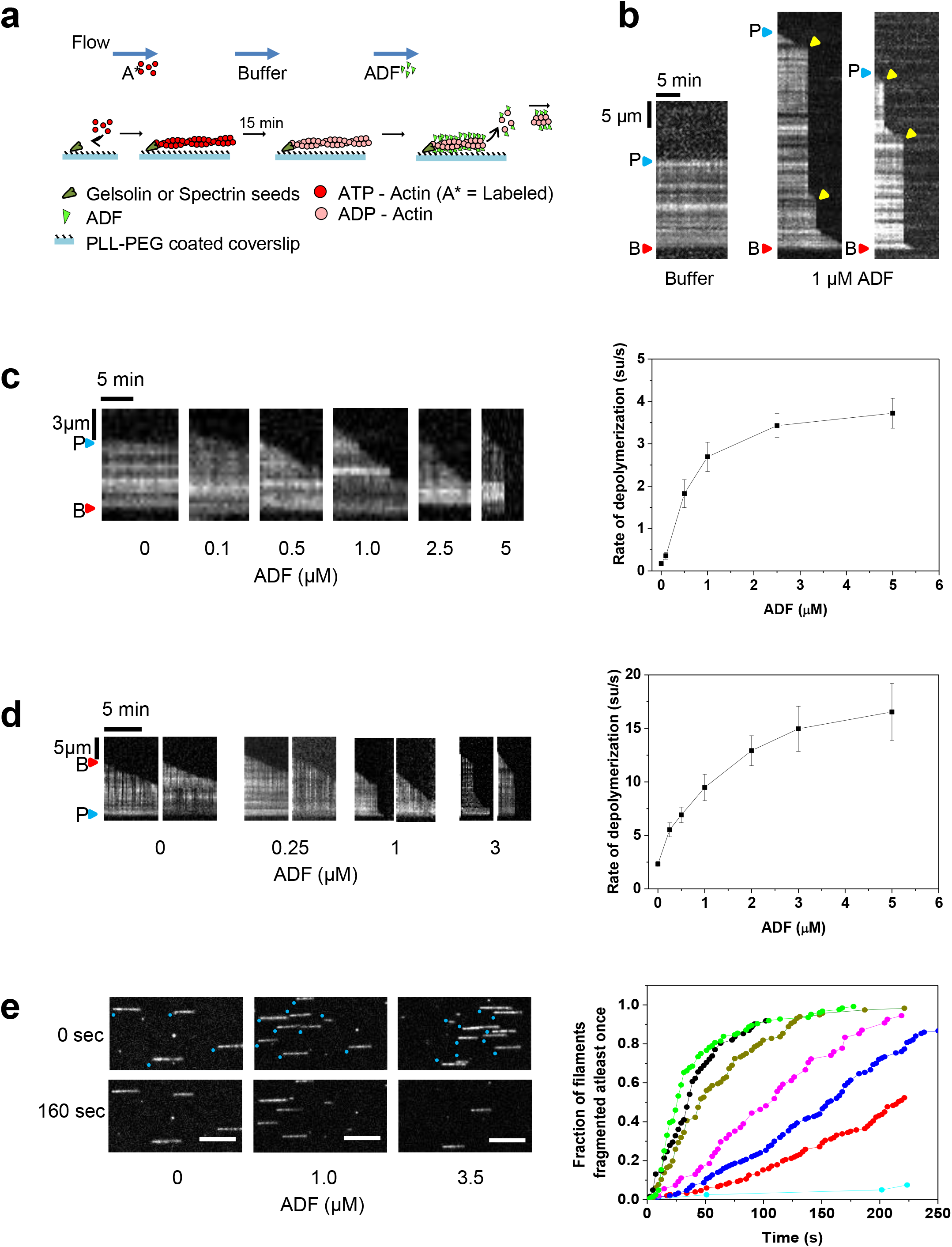
ADF severs filaments and enhances depolymerization at filament pointed and barbed ends. **(a)** Schematics of the set-up for single filament depolymerization. Filaments exposing free pointed ends or barbed ends were grown from coverslip-anchored gelsolin-actin complexes or spectrin-actin seeds respectively, and exposed to F-buffer and ADF in F-buffer. **(b)** Representative kymographs of ADP-F-actin filaments anchored at barbed ends (red triangle) (B) and exposing free pointed ends (P) (blue triangle) to either buffer only (left panel) or 1 μM ADF in buffer (2 right panels). Severing events: yellow triangles **(c)** ADF concentration dependence of pointed-end depolymerization. Left: Representative kymographs. Right: Rate of pointed-end depolymerization versus ADF. **(d)** ADF enhances depolymerization at filament barbed ends. Left: Representative kymographs. Right: Barbed-end depolymerization rate versus ADF concentration. Error bars: S.D. each data point represents the average of 50 measurements cumulated from 3 independent experiments. **(e)** Time-lapsed images of ADP-F-Actin filaments with their pointed ends anchored (blue dots) and barbed ends capped being severed by ADF (flow direction: left to right) (left) and Cumulative distribution function of the first severing event per filament as a function of time for different ADF concentrations – No ADF (cyan), 0.5 μM (red), 1 μM (blue), 2 μM (magenta), 3.5 μM (dark green), 5 μM (black) and 10 μM (light green). Each curve consists of about 70-80 filaments of average length around 10μm.

To evaluate the effect of ADF on depolymerization at barbed ends, filaments were grown from coverslip-anchored spectrin-actin seeds (See Figure 1a for schematics). These filaments were aged to ADP-F-actin by exposing them to F-buffer for 15 minutes to enable phosphate release. The free barbed ends of these filaments were then exposed to ADF in F-buffer as above. In buffer alone, barbed ends depolymerize at 2.31±0.27 su/s, similar to previous reports of ∼ 1.4 su/s [33, 35]. The barbed-end depolymerization rate increased with ADF in a saturating fashion (Figure 1d) up to about 16.54±2.68 su/s at 5μM ADF (n=50 filaments, 3 independent experiments), in a trend similar to that seen for pointed ends (Figure 1c). ADF’s destabilization of actin-actin interactions in the filament [36] which promotes filament severing [17–19, 37] might also cause the so far not detected enhanced disassembly at filaments ends. Why hasn’t this been seen earlier? A possible explanation lies in the methods used, and maybe in the acknowledged quantitative differences between the various cofilins and ADF variants ([38, 39]. In previous studies, filaments were mostly maintained in the evanescent field by surface anchoring [17, 18]. The filament fragments generated by severing immediately left the evanescent field [18], while shortening of the anchored filaments was interpreted as disassembly [20]. In our method, the flow maintains filaments aligned in the evanescent field, which allows visualization and accurate measurement of both severing and depolymerization. Future experiments with fluorescently labeled ADF will be valuable in confirming the results presented here. However, this has so far been a challenge because it has 8 cysteine residues, many of which are close to actin-interaction regions [18].

Recently Twinfilin and Srv2 were together found to promote a similar increase in pointed-end (17-fold) and barbed-end (3-fold) depolymerization [35]. Aip1 also enhances depolymerization of ADF/Cofilin-decorated filaments [29, 40]. However, enhanced end depolymerization has so far never been directly visualized for ADF/Cofilin alone. Our results demonstrate that ADF alone is sufficient for both fragmentation and depolymerization of filaments. The severing activity of Cofilin has been reported to decrease at high cofilin concentrations [17]. Severing appeared optimal at the boundary between ADF-decorated and bare regions of the filament [19]. Results presented here, show that both filament severing and depolymerization increase with ADF concentration up to about 5 μM. Both properties reflect the lower structural [36], mechanical [41] and thermodynamic stability of ADF-decorated than bare filaments, testified by the higher critical concentration for assembly of ADF-F-actin than F-actin in ADP [21]. The severing mechanism used by ADF thus differs from the “hit and cut” mechanism used by gelsolin, Spire or Cobl.

### ADF increases steady-state monomer concentration

As discussed previously [1], severing of pure actin filaments by itself does not change the steady-state monomer concentration (∼0.1 μM). ADF’s effect on G-actin concentration has so far mostly been evaluated by bulk solution assays - SDS-PAGE of supernatants of high speed centrifuged F-actin, or measurements of sequestered ATP-G-actin in the presence of Thymosin β4 [42]. Nevertheless, it can be argued that very short filaments may not be pelleted by highspeed sedimentation, thus causing an over-estimation of monomeric actin in the supernatant.

We therefore designed a novel microfluidics-assisted TIRFM assay to directly measure the monomeric actin concentration and monitor how ADF affects it. A mix containing 5 μM 10% Alexa 488 labelled F-actin, 3μM Profilin and varying amounts of ADF was prepared in F-buffer and allowed to incubate to reach steady-state (∼2 hours, [21]). Tracer filaments exposing free barbed ends were then elongated from immobilized spectrin-actin seeds in a flow containing 1 μM Profilin-Actin (10% Alexa 488 labelled actin) (see Figure 2A for a schematic). These filaments were then exposed to a flow containing the steady-state mix with F-actin, ADF and Profilin. The monitored filaments acted as” tracers” reporting the actual assembly dynamics of filament barbed ends in the population of flowing filaments. The elongation rate of the tracer filaments thus is a reliable measure of the concentration of actin monomers in the steady-state flow. This experiment was repeated at different ADF concentrations to observe the effect of ADF on steady-state concentration of monomers co-existing with filaments. (Figure 2a).

**Figure 2:**
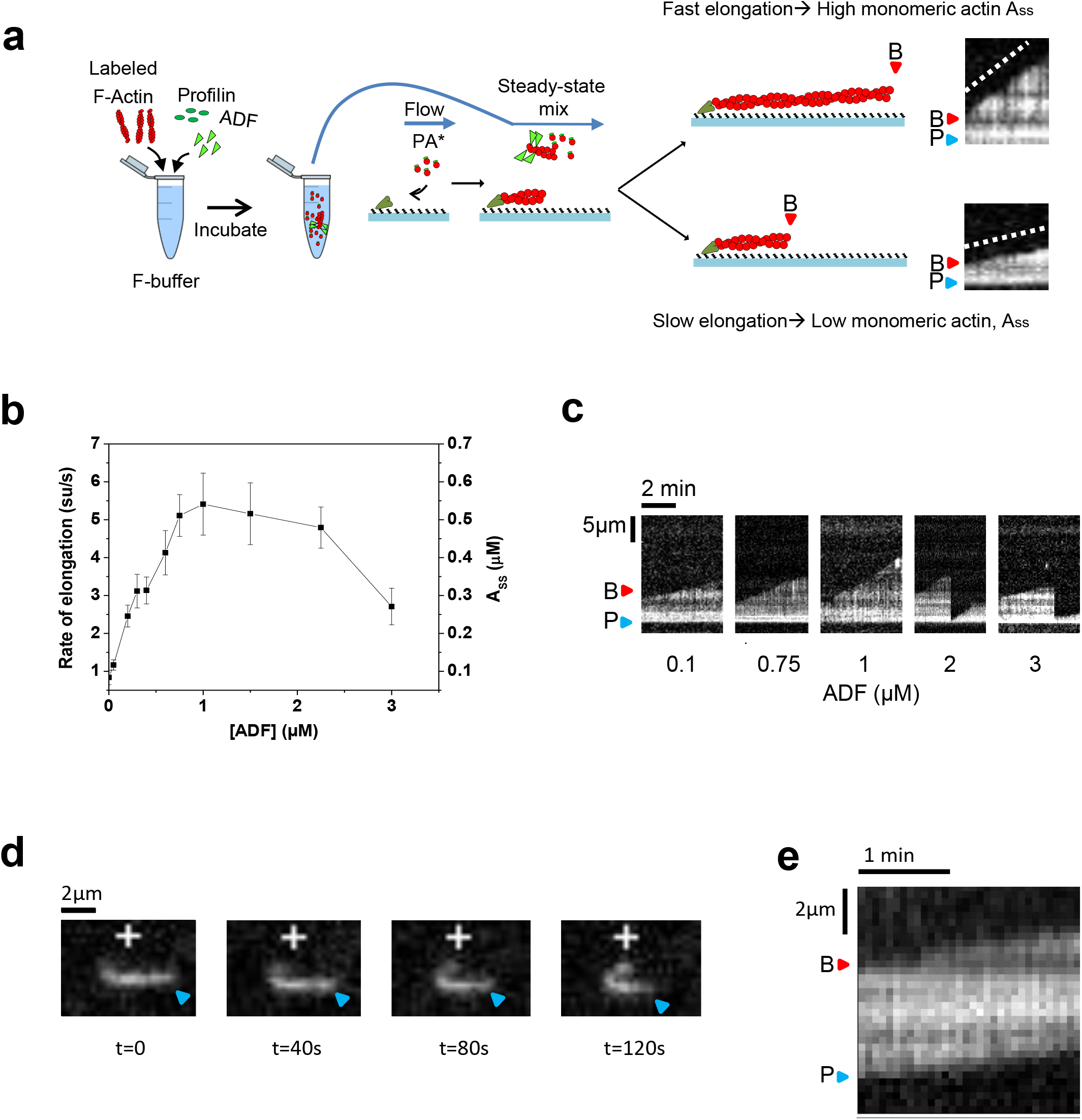
ADF increases the steady-state concentration of monomeric actin (Ass). **(a)** Sketch of the setup to measure the amount of steady-state ATP-monomers as a function of ADF. Tracer filaments expose free barbed ends to solutions of pre-assembled F-actin, 3 μM profilin and varying amounts of ADF at steady state. Example kymographs show fast/slow elongation as indicators of high/low values of the steady state amount of profilin-bound actin-monomers (PA_ss_). (Pointed ends: P, blue triangles; Barbed ends: B, red triangles) **(b)** Elongation rate of tracer barbed ends as a function of ADF concentration (Error bars: S.D. Each data point averaged from at least 50 filaments over 3 experiments). **(c)** Representative kymographs. **(d)** Filaments from the flow (0.75 μM ADF), incidentally stuck to the coverslip surface, display barbed end (+) elongation and pointed end shrinkage. **(e)** Kymographs showing constant filament length maintained during treadmilling. B and P indicate the two ends.

In absence of ADF, tracer barbed ends grew very slowly (0.14±0.05 su/s, equal to the rate of pointed-end disassembly). In the presence of ADF, tracer barbed ends grew faster, up to about 4 su/s at 2 μM ADF, which translates into a concentration of 0.4 μM profilin-actin at steady state (Figure 2b). The decline observed at higher ADF likely reflects ADF-mediated inhibition of conversion from ADP-actin to ATP-actin monomers [21]. ADF-induced severing events are visible between periods of barbed-end growth (Figure 2c). Interestingly, a few filaments from the bulk flow sometimes landed and bound non-specifically to the glass surface, exposing both of their two ends to the flow. Since these filaments were part of the bulk flow (unlike the tracer filaments), they were actually at steady state. These filaments undergo barbed-end growth and pointed-end shrinkage at identical rates, maintaining constant length, i.e. treadmilling (Figure 2d, 2e). In conclusion, ADF promotes the establishment of a higher steady-state concentration of ATP-G-actin associated with enhanced treadmilling.

### Capping protein funnels monomers to uncapped barbed ends

Capping of barbed ends is also essential for actin-based motility [11, 24, 26]. By blocking a large fraction of filament barbed ends, Capping Protein has been proposed to “funnel” actin monomers released from all pointed ends onto a few non-capped filaments, locally initiated in cells by surface-bound nucleators, which thus grow faster [10, 11]. Funneling is expected to be most effective between 90% and 99.9% capping, where steady-state monomer concentration would steeply increase [43]. Pending direct visualization of this funneling behavior, this hypothesis has been challenged [16]. We therefore investigated the “funneling effect” of CP using the above assay.

Tracer filaments exposing free barbed ends were now exposed to a solution of F-actin preassembled at steady state with profilin (3 μM) and varying amounts of CP. Profilin prevents monomer binding to pointed ends. At a CP concentration of 0.2 nM, at which 67% of bulk filaments are capped, the tracer barbed ends elongate at 0.53 ± 0.14 su/s, corresponding to a concentration of ATP-actin monomers of 150nM (Figure 3a, b). At 5nM CP (98% capping of bulk filaments) the growth rate of 8.06 ± 0.28 su/s corresponds to a steady-state concentration of ATP-monomers (essentially profilin-actin) of 0.8 μM. Tracer ends also get capped at a rate increasing with CP concentrations (k_+C_ = 4.06±0.17 μM^−1^s^−1^) (Figure S1), in agreement with earlier studies [44]. The data agree with the non-linear dependence of the steady state monomer concentration on the extent of capping (Figure 3b) [27, 43].

**Figure 3:**
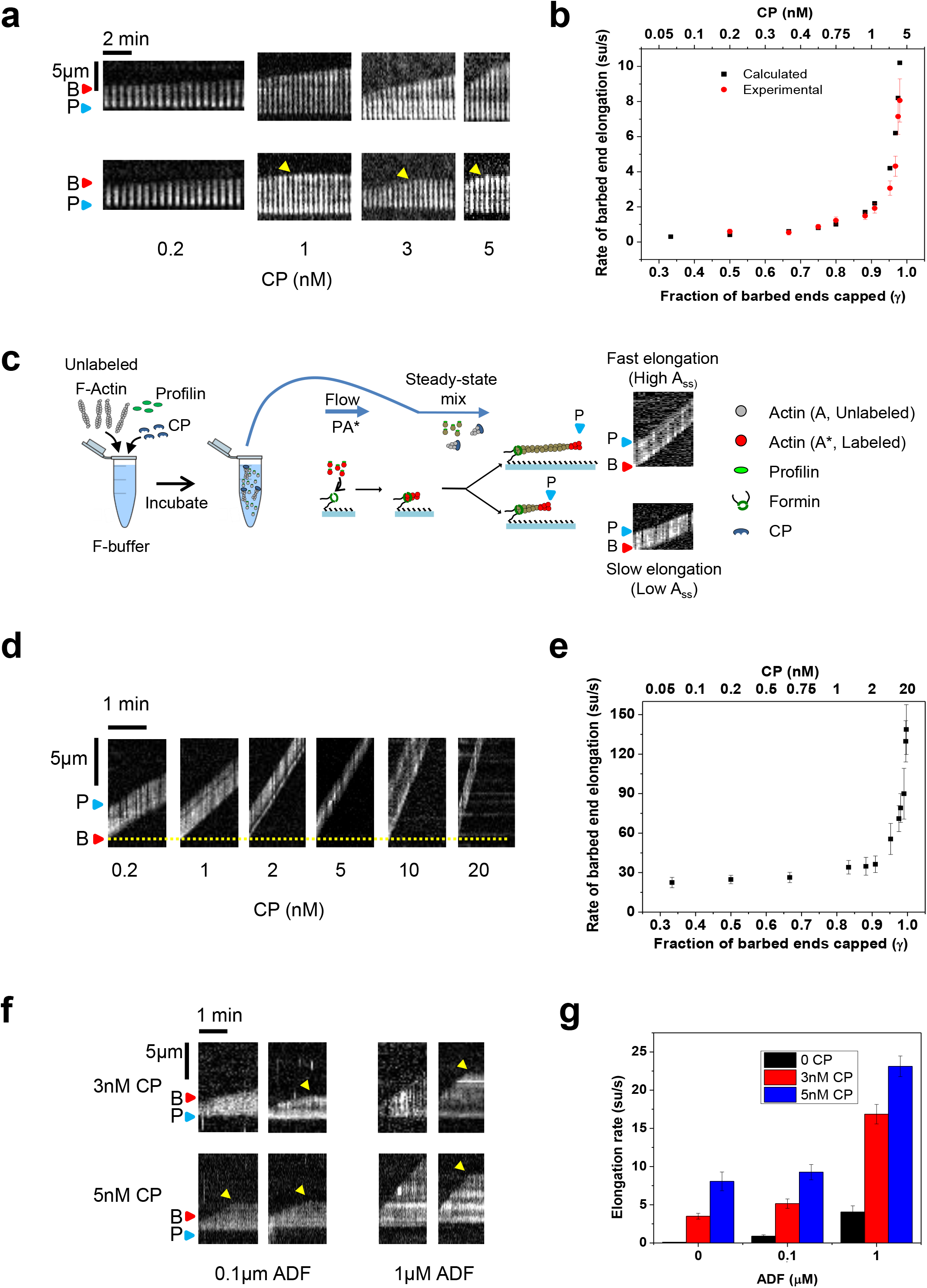
Capping protein increases barbed-end elongation rate in synergy with ADF. Experimental conditions are as in Figure 2 except ADF replaced by CP. Tracer barbed ends were exposed to a flow containing preassembled 5 μM F-actin (Alexa 488 labelled), 3 μM profilin and CP. **(a)** Representative kymographs of free barbed ends elongating (2 for each condition). Arrest of growth by capping is, indicated by yellow arrows. (Pointed ends: P, blue triangles; Barbed ends: B, red triangles) **(b)** Rate of barbed-end elongation versus CP concentration: experimental and calculated (See Sup. Info) **(c)** Sketch of the setup to measure the amount of steady-state ATP-monomers as a function of CP concentration using formin-bound barbed ends. Tracer filaments are nucleated by exposing anchored formins to profilin and labeled actin (Alexa 488). These tracers are then exposed to steady state solution containing unlabeled actin. The labelled segment of the filament appears to move away as unlabeled monomers are added by the formin at point of anchoring. Example kymographs show fast/slow elongation as indicators of high/low values of the steady state amount of profilin-bound actin-monomers (PAss) **(d)** Representative kymographs of barbed ends elongating from anchored formins (yellow dotted line). F-actin in the flow was unlabeled, letting the labelled pointed-end region of tracer filaments move away as insertional polymerization occurs at formin-bound barbed end. **(e)** Rate of formin elongation versus CP concentration. **(f)** Representative kymographs (2 for each condition) of tracer barbed ends growing in the presence of CP and ADF. **(g)** Diagram representing the synergy between CP and ADF in enhancing steady state growth of individual filaments. Error bars: S.D. Each data point averaged from at least 50 filaments over 3 experiments.

At CP concentrations above 5 nM, tracer filaments get capped too rapidly for their elongation rates to be measured. We therefore used formin-bound barbed ends as tracers, because CP binds to formin-bound barbed ends with a 500-fold lower affinity than to free barbed ends [45, 46], allowing us to test higher CP concentrations up to 20 nM (99.5% capping of the flowing filaments). Fluorescent filaments were nucleated from surface-anchored formin mDia1 molecules by exposing them to a flow containing 1 μM labelled G-actin and 1 μM Profilin. These formin-elongating “tracer” filaments were then exposed to a flow containing unlabeled 5 μM F-actin, 3 μM Profilin and varying amounts of CP. Due to the unlabeled nature of actin monomers generated in the steady-state flow, fluorescent fragment of the tracer filament appears to move as monomers are insertionally added at the site of formin anchoring (Figure 3c). The rate of formin-bound barbed-end assembly of tracer filaments increases with CP up to 130 su/s at 20 nM CP (Figure 3d,e). This value translates into a steady state concentration of profilin-actin of 2.6 μM, assuming a value of 50 μM^−1^.s^−1^ for the association rate constant of profilin-actin to formin-bound barbed ends [32]. This value indicates that 87% of the 3 μM profilin is bound to actin, consistent with about 100% capping. In conclusion, capping of majority of bulk filaments at steady state enhances the growth rate of a few uncapped (tracer) filaments due to an increase in polymerizable monomers, thus demonstrating the existence of the “funneling effect”.

In providing single filament evidence for the funneling effect of Capping Protein, our data clarify a discrepancy between the reported enhancement of actin-based propulsion of *Listeria* and N-WASP coated beads by CP and its inhibition at high capper concentrations [10, 25], and the different observations and proposed interpretation of the function of CP made by Akin and Mullins [16]. A key difference between the experiments of Akin and Mullins and those presented here is that while the flowing actin solutions in the present study consist of F-actin (and actin monomers) at steady state, Akin and Mullins conducted experiments that contained only G-actin at time zero. Hence the rate of actin assembly and resulting propulsion rates varied in a non-defined fashion as the G-actin was depleted with time. A recent theoretical study argued that CP only locally increases monomer concentration near the location of branched nucleation [15] (e.g. near an ActA coated bead), in contrast with our results showing a global bulk increase in monomer concentration. Unfortunately, it is not possible to quantify the rate of nucleation in our study. In conclusion, our *in vitro* results obtained at steady state conditions seem to account for *in vivo* effects of CP on motility [1].

### ADF and Capping Proteins synergistically enhance the concentration of polymerizable actin monomers and barbed-end assembly at steady state

We have thus far shown that ADF and CP individually increase the concentration of polymerizable actin monomers co-existing with filaments. We next explored how they functionally cooperate when present together in F-actin solutions. Tracer filament barbed ends were exposed to F-actin solutions at steady state, containing either ADF alone, CP alone or both (Figure 3f,g). CP and ADF appear to synergize in eliciting even faster barbed-end growth rates than individually. At 1 μM ADF and 5nM CP, the elongation rate of tracer barbed ends was 23 su/s as compared to 4 su/s in presence of 1 μM ADF alone and 8 su/s in presence of 5nM CP alone. ADF and CP therefore combine to enhance the elongation rate 160-fold (∼0.14 su/s in their absence), as compared to in absence of both CP and ADF.

In the presence of ADF, the concentration of ATP-monomers is established at a higher value to ensure equal global (bulk) rates of barbed-end growth and pointed-end depolymerization (Figure 4c). This effect cumulates with CP-mediated funneling (Figure 4b, e), in a synergistic fashion (Figure 4d, e). The individual and cooperative regulatory activities of ADF and CP have been demonstrated here at the single filament scale, thanks to the original approach used for direct TIRF-based measurement of the concentration of polymerizable monomers. The values of the measured growth rates are consistent with the ones expected from a graphical representation of the contribution of barbed and pointed ends of filaments in the steady state of actin assembly (Figure 4e). Elongation rates measured here (23 su/s) at 1μM ADF and 5 nM CP are in close to barbed-end assembly rates estimated in the lamellipodium (66 su/s) [47, 48]. The results presented here represent a step forward in our understanding of the molecular mechanisms of how cells might control a steady-state monomer pool that defines the rate assembly at barbed ends. Our method can easily be adapted to integrate the mechanistic contributions of other effector proteins (e.g. Aip1, Twinfilin, Coronin) regulating actin treadmilling in cells. Thus it promises to be seminal in quantitatively addressing, at the single filament level, questions such as the actual concentrations of profilin-actin in various cell extracts, which is a major unsolved issue [1, 49, 50].

**Figure 4:**
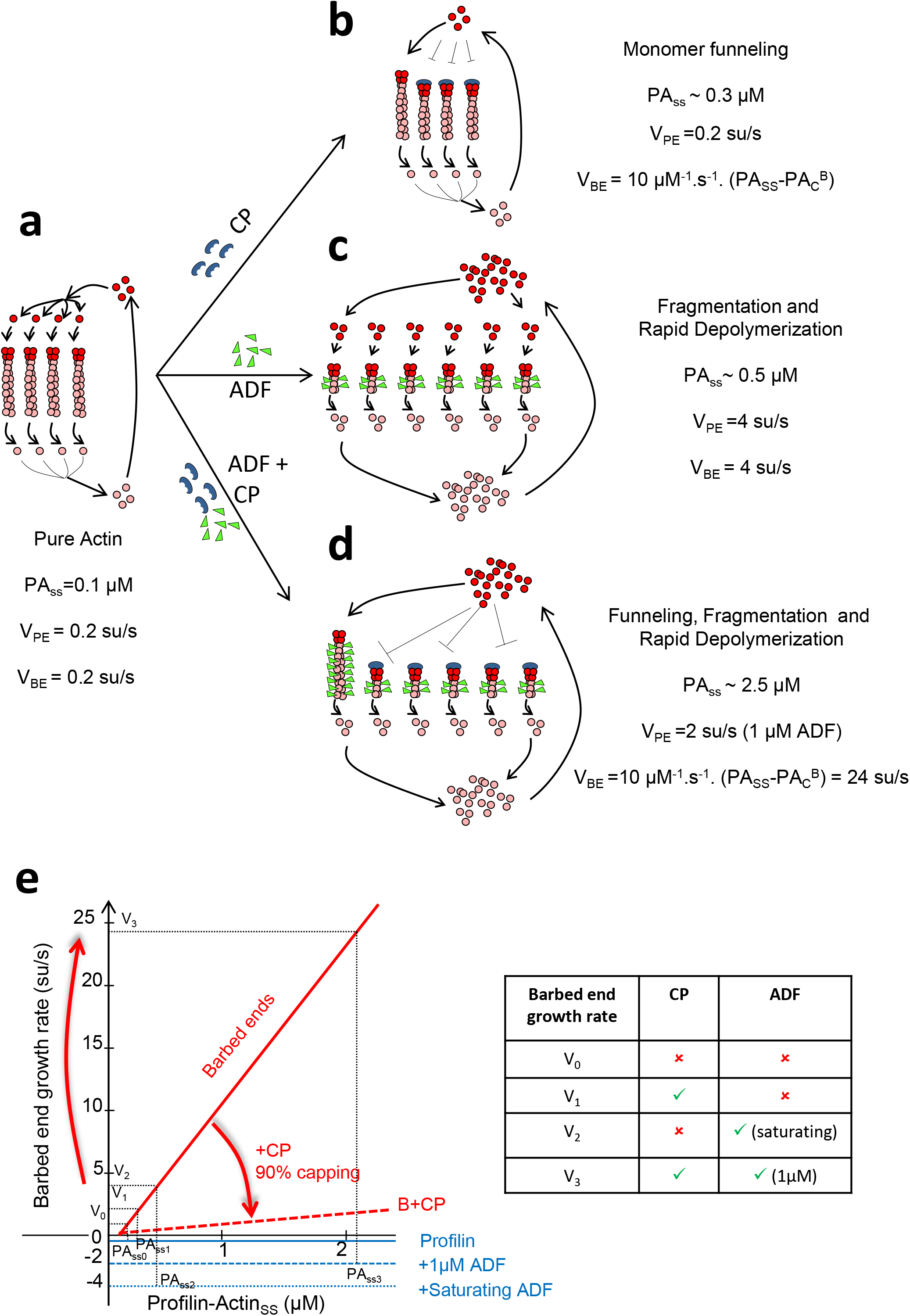
Combined effects of ADF and CP on barbed-end growth rate at steady state. **(a)** Control filaments (4 shown) treadmill very slowly due to low steady-state profilin-actin concentration (PASS = 0.1 μM) at which barbed-end growth rate (V_BE_= k+_B_.(Ass.C_c_^B^) balances slow pointed-end depolymerization (Vpe∼0.2 su/s) **(b)** In the presence of CP (90% capping), all pointed ends feed a few much faster growing uncapped barbed ends. **(c)** In the presence of ADF, enhanced pointed-end depolymerization (4 su/s at saturation by ADF) establishes a higher concentration PASS (0.5 μM). **(d)** In the presence of both ADF and capping protein (CP), PASS further increases and non-capped barbed ends grow faster. **(e)** Graphical representation of the effects of ADF and CP on the contribution of barbed and pointed end dynamics in establishing the steady state of actin assembly. Red and blue lines represent the change in elongation rates at filament barbed and pointed ends versus monomer concentration, in the presence of 3 μM profilin. Pointed ends only depolymerize at constant rate, 0.2 su/s and 2 su/s in absence (continuous line) or presence (dashed line) of 1 μM ADF. Barbed ends grow with 100% free (non-capped) barbed ends (continuous red line) or 90% capped barbed ends (dotted red line). The steady state monomer concentration is first determined as the monomer concentration at which pointed end disassembly equals barbed end assembly bulk rate. The elongation rate V_i_ of individual barbed ends at steady state is then read on the graph as the rate of non-capped barbed end at the steady state monomer concentration. Conditions: no regulator (V_0_); + 2 nM CP: (V_1_); + 1μM ADF (V_2_); + both ADF and CP (V_3_).

## Author Contributions

SS and MFC conceived and designed the experiments, SS conducted the experiments and analyzed the data, SS and MFC wrote the manuscript.

## Acknowledgements

We thank Bruce Goode, Jane Kondev, Lishibanya Mohapatra Adam Johnston, Silvia Jansen, Greg Hoeprich, for stimulating discussions on actin disassembly and for critical feedback. We thank Julien Pernier for assistance in expression and purification of ADF and Bérengère Guichard for technical assistance in expression and purification of all other proteins. The work was supported by an ERC advanced grant “Forcefulactin” # 249982 and a EC FP7 grant # 241548 to M.-F. C.

